# Frontal theta power prospectively associated with response inhibition

**DOI:** 10.1101/2024.05.13.593803

**Authors:** Daan van Rooij, Sam van Bijnen, Iris Schutte, Nathan van der Stoep, J. Leon Kenemans

**Affiliations:** Experimental Psychology, Helmholtz Institute, Utrecht University, Netherlands

**Keywords:** Response inhibition, proactive inhibition, stop-signal performance, frontal theta power

## Abstract

A proactive mechanism has been postulated to promote successful inhibition (Kenemans, 2015). Specifically, this mechanism is thought to operate before any action demanding or countermanding event has occurred. In the current study, we investigated whether EEG theta power could reflect this mechanism, in a sample of healthy individuals performing a stop-signal paradigm. By comparing frontal theta power preceding failed versus successful stop trials, we tested whether frontal theta is predictive of inhibition success. We hypothesized that proactive cognitive control manifests in frontal theta power preceding a countermanding go-stop event. Our results demonstrate that frontal theta is indeed higher preceding successful as compared to preceding failed stopping events. We also show that frontal theta power preceding stopping events is associated with Stop-Signal Reaction Times (SSRT), with a higher theta being indicative of shorter SSRTs. This association was not present for go-RT. This study may be the first to reveal a relationship between lower frontal theta power and subsequent stopping failure, suggesting theta’s role in proactive response inhibition.

## Introduction

Cognitive control is an umbrella term referring to a collection of functions such as attention, working memory, conflict monitoring, and response inhibition. Response inhibition is an important mechanism for suppressing a prepared or prepotent response tendency. Previously, we postulated that response inhibition involves two components: a proactive cognitive-control component and a reactive component. The reactive component refers to the ability to process and react to an unexpected event, whilst the proactive component serves to prepare the neural pathways that facilitate the reactive (or another) component, should response suppression prove necessary (Kenemans, 2015; see also Cai et al., 2011 and Majid et al., 2013). Response inhibition is preferably assessed using a stop-signal task (SST, e.g. Verbruggen et al., 2019), in which subjects respond to various go-stimuli which are occasionally followed within a split-second by a stop-signal, signaling the subjects to suppress the ongoing response tendency. From a stop-signal task, a stop-signal reaction time (SSRT) can be derived, which reflects the time needed to process the stop signal and suppress the response that was prepared.

Theta oscillations as derived from the EEG over frontal cortical regions have been identified as reflecting a pivotal mechanism underlying cognitive control (Cavanagh & Frank, 2014; Zavala et al., 2018). However, it is unclear whether frontal theta also reflects cognitive control in the context of response inhibition. Several previous findings indicate that theta power might be associated with proactive cognitive control. For example, increased theta power has been shown to be elicited by cues that inform about an upcoming event (as compared to non-informative cues; Mazaheri et al., 2010) or the most likely response to an event (Mazaheri et al., 2014), as well as by cues that signal highly probable subsequent response conflict (Van Driel et al., 2013). The same holds for cues that signal task switching. Such cues elicit stronger theta connectivity (Cooper et al., 2015), or theta power (Cooper et al., 2017), relative to repeat-task cues. In contrast, one study did not observe differences in pre-trial theta as a function of nogo-probability in a go/nogo task (Adelhofer et al., 2020). However, as noted by Aron (2011), in a go/no-go paradigm one cannot be sure when the Go and Stop processes begin. As a result, the timing of any proactive response-inhibition mechanism becomes uncertain.

If a proactive component does exist for response inhibition in a stop-signal task, it could manifest as enhanced frontal theta power immediately preceding go stimuli in a stop-signal context. Theta power would then reflect the process of cognitive preparation for a potential inhibition event. Following Kenemans (2015): The reactive component of response inhibition would be implemented as a frontal (f) P3 stop effect (i.e. the difference in fP3 to the stop signal between successful and failed stops), most likely implemented in the pre-supplementary motor area (SMA). The proactive component should be visible as the modulation of stop success in response to the stop signal, which is thought to reflect proactive preparation in prefrontal regions (e.g., the right inferior frontal gyrus; Aron et al., 2006). Therefore, we expect to observe increased theta power measured above the prefrontal areas before trials where a subject succeeds in stopping a response as compared to those where a subject failed. To the best of our knowledge, the association between frontal theta and stopping performance has not been investigated in an SST paradigm. Furthermore, it is important to dissociate proactive inhibition from proactive slowing. It has been reliably demonstrated that in SST paradigms, participants often show proactive slowing of their reaction to the go-stimulus in anticipation of an expected stop event (Vink et al., 2014; Zandbelt et al., 2013; Pas et al., 2017; Pas et al., 2021). This can be observed as longer reaction times with higher stop-signal probability. However, an increase in proactive inhibitory control would be associated with faster inhibition and hence shorter SSRT but not with longer reaction times on go-trials (go-RT). This would constitute a yet further instance of dissociations between SSRT and go-RT, such as repeatedly demonstrated for stop-signal probability which affects go-RT but not SSRT (Kenemans, 2015; Lansbergen et al., 2007).

Hence, in the current study, we investigated EEG theta power in a sample of healthy individuals performing a stop-signal paradigm. In this paradigm, we looked at the stop-trials, which are 25% of trials where a go-signal is followed after a split-second by a stop signal. By comparing frontal theta power preceding failed versus successful stop trials, we aim to test whether frontal theta is predictive of inhibition success. We hypothesized that proactive cognitive control manifests in frontal theta power preceding a countermanding go-stop event. Thus, we expect higher frontal theta preceding successful vs failed stop events. Additionally, if frontal theta power is specifically indicative of increased proactive control, we expect overall theta power preceding go-stop events to be associated with shorter average SSRT values but not with longer go-RTs. Lastly, we will compare two modalities for the stop signal: a visual and an auditory stop signal variant, to rule out that theta power and inhibition success are dependent on the sensory modality of the stop signal. We expected no difference in theta power between these conditions (Kenemans et al., 2023).

## Method

### Participants

EEG and stop-signal task data were acquired for 52 healthy volunteer participants (mean age = 24.2, SD = 4.5; N females = 31). All participants signed an informed consent form and reported normal hearing and normal or corrected-to-normal vision. Participants were paid 20 Euros for their participation. The study was approved by the local ethics committee of the faculty of Social and Behavioral Sciences of Utrecht University.

### Procedure

Participants read and signed the informed consent and provided demographical information before the EEG cap and EEG compatible in-ear headphones (13 or 10mm based on the participant’s ear size, Etymotic research, inc.) were placed on the head. Participants were seated in a dimly-lit room, with their head on a chin-rest 60 cm centered in front of the computer screen. After receiving the instructions, subjects performed the stop-signal task (see details below) for approximately 35 minutes (plus a 5-min break halfway) during which EEG was recorded.

### Stop-signal task

The stop-signal task (SST) consisted of a two-choice response task (Kenemans et al., 2023). Participants had to discriminate between two go-stimuli: the letters “X” and “O” (visual angle for both letters, 1.6° x 1.6°), by pressing the left or the right button with the left or the right index finger, respectively. The mapping of “X” and “O” to either left or right was counterbalanced over blocks and conditions.

A single total trial duration was initialized randomly to last between 1500 and 1518ms, during this entire time a fixation cross remained onscreen. The go stimulus was then presented first, for 150ms, centrally 2cm above the fixation cross. On 25% of the trials, a stop-signal was presented after the go-signal, following a variable Stimulus Onset Asynchrony (SOA, see below for detailed description). For each subject, two conditions were presented based on the nature of the stop-signal. In the visual condition, the stop-signal consisted of a screen with a red background, presented for 150ms. In the auditory condition, the stop-signal consisted of a 1000hz tone (72dB), presented for 150ms in both ears via in-ear headphones. After the go- and potential stop-signals were presented, the Inter-Trial-Interval (ITI, consisting of just the fixation cross) was calculated to fill the remainder of the total trial duration.

The experiment consisted of a total of 13 blocks, of which 5 practice blocks and 8 experimental blocks, divided over 4 task (2 stop-signal modalities x 2 response mappings) conditions. The first practice block consisted of 24 go trials, and was also used to establish a baseline reaction time on go-trials (go-RT).

Next, the 3 blocks per condition were presented (1 further practice block and 2 experimental blocks). The different conditions were blocked and counterbalanced, so that all subjects were presented with both visual and auditory stop-conditions, and the “X” and “O” mapped to left/right hand respectively in a counterbalanced way. Each block consisted of 128 trials. The first block of each condition was a practice block, used to update the average SOA. Practice blocks were not included in the eventual analyses, so only the 8 experimental blocks were analyzed, for a total of 8*128=1024 trials per subject, of which 25%, or 256 trials, were stop trials. Halfway through the experiment participants were offered to take a short 5-minute break.

As mentioned above, the SOA was adjusted during the experimental blocks to adapt to the individual participants performance. If subjects managed fewer than 13 successful stops, they were instructed to please slow down a bit, when they managed more than 19 successful inhibition, they were instructed to please speed up a bit. The SOA between the go and the stop stimulus was set at 250ms at the start of the experiment, and was adjusted using a tracking algorithm during each practice block, by increasing the SOA with 50 ms after each successful stop, and shortened with 50 ms after each failed stop trial, leading to an adapted difficulty for every subject. The resulting final value of each practice block (with a minimum possible SOA of 250ms) was set to be the average SOA for the subsequent experimental block. After each experimental block, the average SOA was further updated by subtracting the observed successful stop rate from the expected 50%, and adjusting the new SOA by 2x the difference in percentage converted to milliseconds (de Jong et al., 1990), with again a minimum value of 250 ms. Within each block, the SOA was subsequently jittered between + and − 99ms around the average over trials (rectangular distribution of jitter values). This method of SOA adjustment should lead to an average successful stop rate of 50% for all participants for the experimental blocks (Verbruggen et al., 2019). If, despite these adaptive mechanisms, subjects still scored outside of the 30-80% successful stop rate for any given block, subsequent calculations of the SSRT can be considered unreliable, and these blocks were excluded from further analyses. Note that the original primary motivation for this jitter approach was to be able to use the Adjar-level-2 procedure to separate go and stop ERPs. However, this is not relevant in the current study where we considered only pre-go theta power.

### EEG data acquisition

EEG signals were recorded with the Active-Two system (Biosemi, Amsterdam, The Netherlands) with 64 Ag-AgCl electrodes. Recording electrodes were placed according to the 10/10 system. Four flat-type EOG electrodes were placed: above and below the left eye and at the outer canthi of both eyes. Offline reference electrodes were placed at the left/right mastoid bones. EEG signals were online referenced to the Common Mode Sense/ Driven Right Leg electrode, sampled at 2048 Hz and online low pass filtered at DC to 400 Hz.

### EEG power analyses

Raw EEG signals were analyzed offline using Brain Vision Analyzer 2.0 (Brain Products GmbH). EEG data were resampled to 64 Hz (after adequate low-pass filtering), re-referenced to the mastoid reference electrodes, and high-pass filtered (0.5 Hz cutoff). Data were divided into 1-s segments (950 ms before to 50 after a go stimulus that was followed by a stop signal). Automatic ocular correction was performed using the EOG channels (Gratton et al., 1983). After ocular correction, automatic artifact rejection was used (maximum allowed absolute difference between two values: 100 μV, lowest allowed activity within a 100 ms interval: 0.5 μV). Trials containing a stop-signal were split for successful and for failed stops resulting in two conditions for the visual stop signal and two conditions for the auditory stop signal.

To validly compare theta power preceding both failed and successful stop-trials, an equal number of failed and successful stop trials has to be included in the analyses. Out of the condition with the highest number of trials, the redundant number of trials were randomly selected and removed. This led to the inclusion of an average number of trials across subjects of between 116 (SD = 13.4) of both failed- and successful stop events.

Subsequently, spectral periodograms were estimated (per trial per condition per participant), using a fast Fourier transformation (Hanning window: 10%) and averaged across trials per condition to obtain power spectra, from which theta-power values were extracted (average power across 3 to 8 Hz).

### Statistical data analyses

To compare the differences in theta power preceding failed vs successful stop trials in both the auditory and visual conditions of the SST, a two-factor within-subject ANOVA was performed, including task performance (failed/successful) and modality condition (visual/auditory) as within-subject factors.

The stop-signal response times were analyzed following the procedure outlined by Verbruggen et al. (2019). First, per block, the proportion of successful stops was calculated. Subsequently, all go-RTs and omissions (for which RT was set at a fixed value of 1500ms) were rank-ordered from shortest to longest. Then the inverse stop rate (= percentage failed stops) was used to identify the corresponding sorted-RT-distribution percentile (e.g., with an inverse stop rate of 45% and 120 sorted go-RTs, the (0.45*120)th go RT was taken). This value was corrected for the average SOA for that block to estimate SSRT for that block. Subsequentially SSRTs were averaged across blocks.

To analyze the relation between relative theta values in the different conditions and stop-signal task performance, a multiple linear regression model was performed including theta power (averaged across failed and successful) as the dependent variable, mean SSRT and go-RT as numerical predictor variables and condition (Auditory/Visual) as within-subject factor.

### Replication sample

To replicate the main analyses of theta power preceding failed and successful inhibition events in a stop-signal paradigm, we additionally included the auditory condition of an existing pilot sample previously described in Kenemans et al. (2023). This sample included a further 28 subjects for which the same SST paradigm (total of 3 blocks with 128 trials each) was acquired using comparable EEG setup and analyses. Statistical data analyses for the replication sample consist of a paired samples t-test comparing the average theta power preceding failed vs successful stop events.

As an additional exploratory analysis, stop-events for the replication sample were separated based on the length of their SOA, grouping all the trials with below-median SOAs (short) and above-median SOAs (long). This allowed us to separate trials based on their difficulty, as long SOA trials are more challenging for the subject to successfully suppress their response. As a secondary analysis, we compared successful stop trials with a long SOA (difficult but successful) with failed stop trials with a short SOA (easy but failed), using a paired-samples t-test. We expected that if frontal theta is reflective of proactive inhibition, the difference between failed and successful trials should be amplified in this secondary analysis.

## Results

Group averages of theta power scalp distribution are depicted in Figure 2.

**Figure 1.**
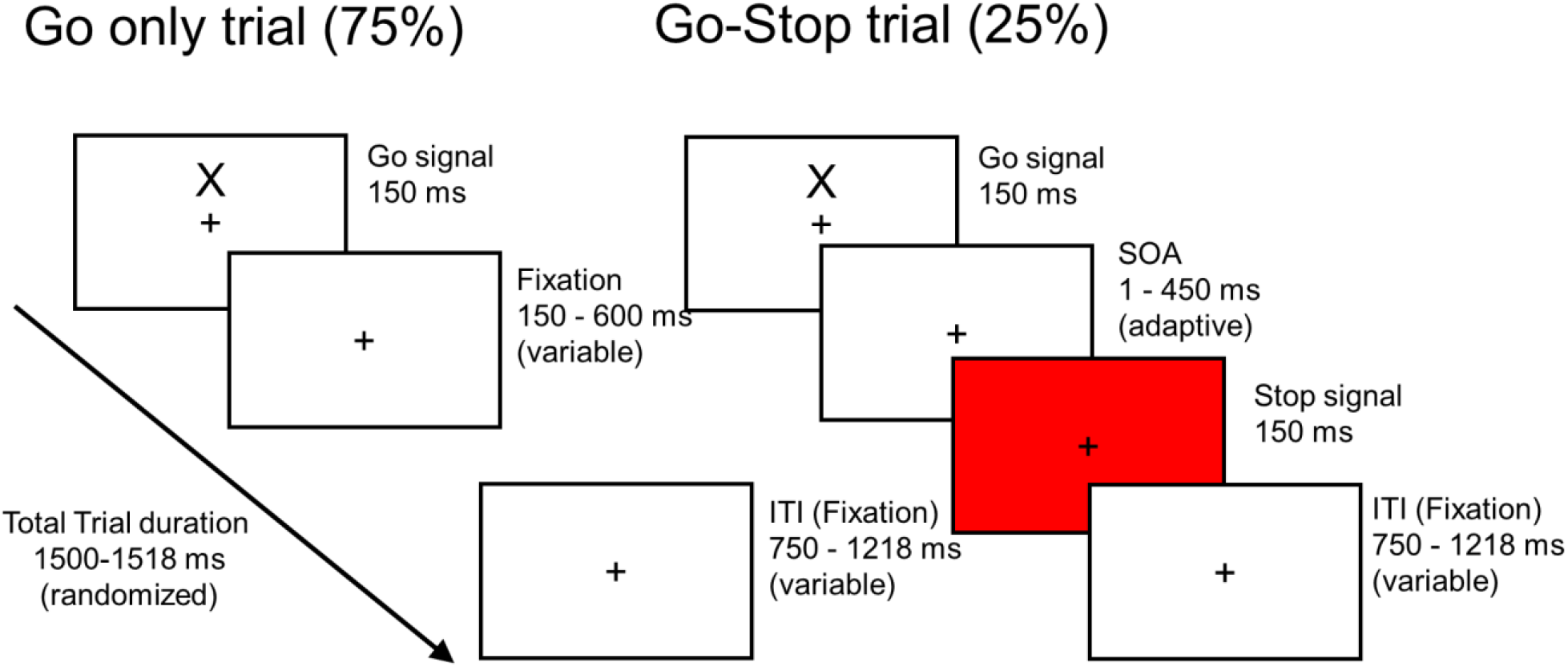
An example of a go (left) and a stop trial with a visual stop signal (right).

**Figure 2.**
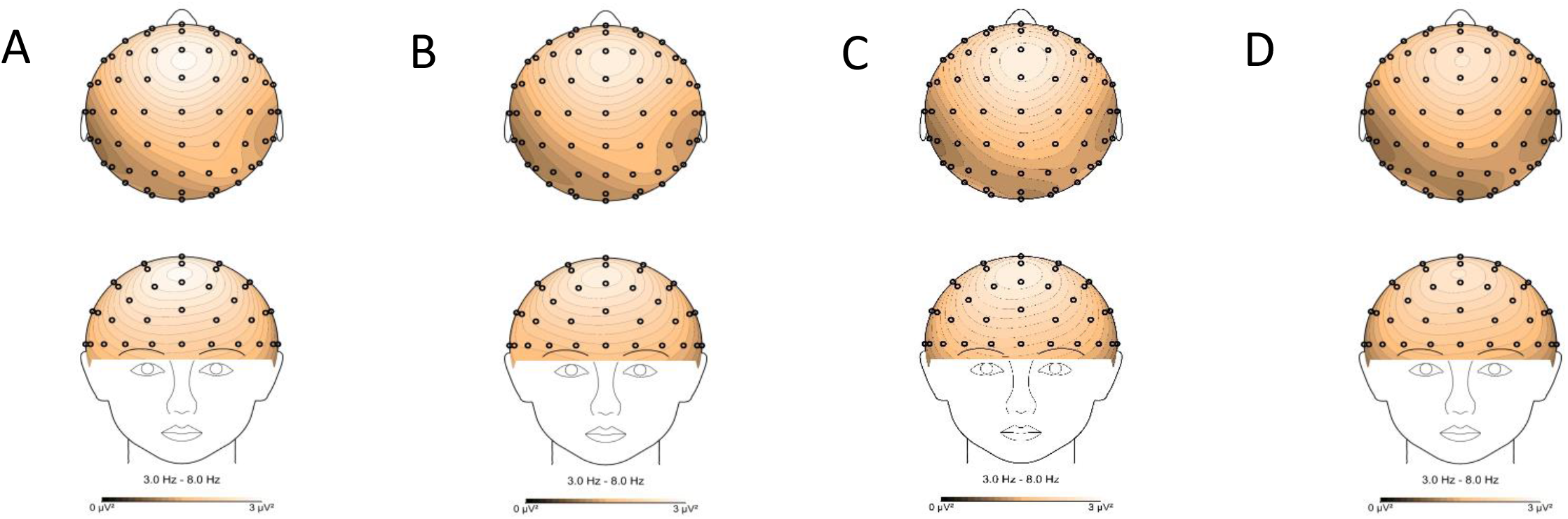
Topography of theta power preceding stop trials separately for auditory successful and failed (A, B), and visual successful and failed (C, D) conditions of the stop-signal task.

### Theta power preceding failed vs successful stop trials

After removing blocks with a stopping performance outside of the accepted range (30-80%, see methods), a total of 5 subjects were removed from further analyses due to too few data points remaining in one or more condition. This led to the final inclusion of 47 subjects in the statistical analyses.

As can be seen in Figure 2, theta power was maximal at the Fz electrode in all conditions. Therefore, for subsequent analyses only Fz theta-power values are used. The EEG power over frequencies from 1-15hz on the Fz electrode for all failed and successful stop trials are depicted in Figure 3.The mean values for Fz-theta-power for failed and successful stop trials in auditory and visual conditions of the stop-signal task are depicted in Table 1 and Figure 4.

**Table 1.** mean frontal theta power values preceding both the Failed and successful trials for both auditory and visual conditions.

| Failed/successful | Auditory/visual | Mean | SD |
| --- | --- | --- | --- |
| failed | auditory | 2.43 | 0.98 |
|  | visual | 2.35 | 0.89 |
| successful | auditory | 2.55 | 1.01 |
|  | visual | 2.47 | 0.96 |

**Figure 3.**
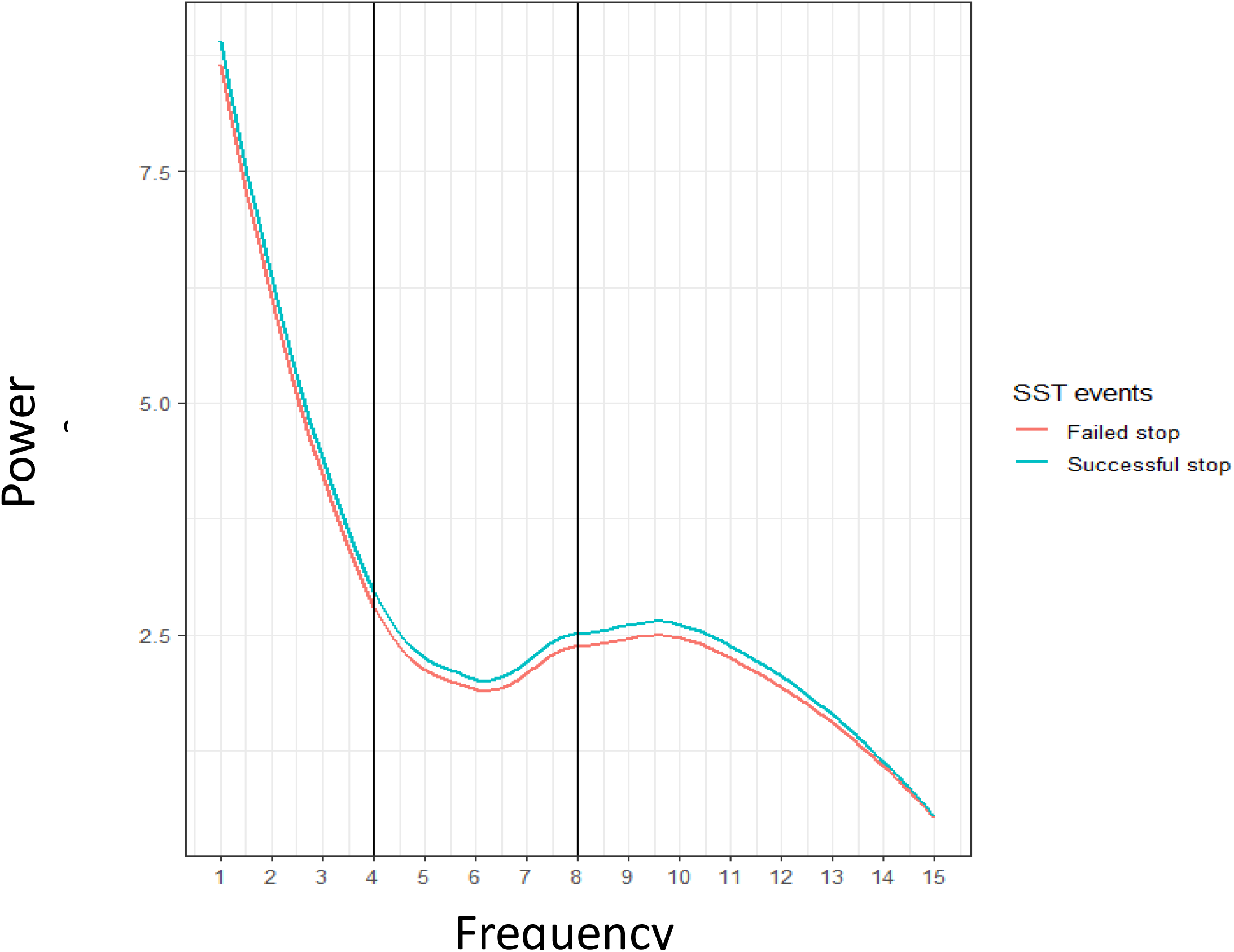
Power spectra over frequencies from 1-15Hz for the second preceding both failed and successful stop events.

**Figure 4.**
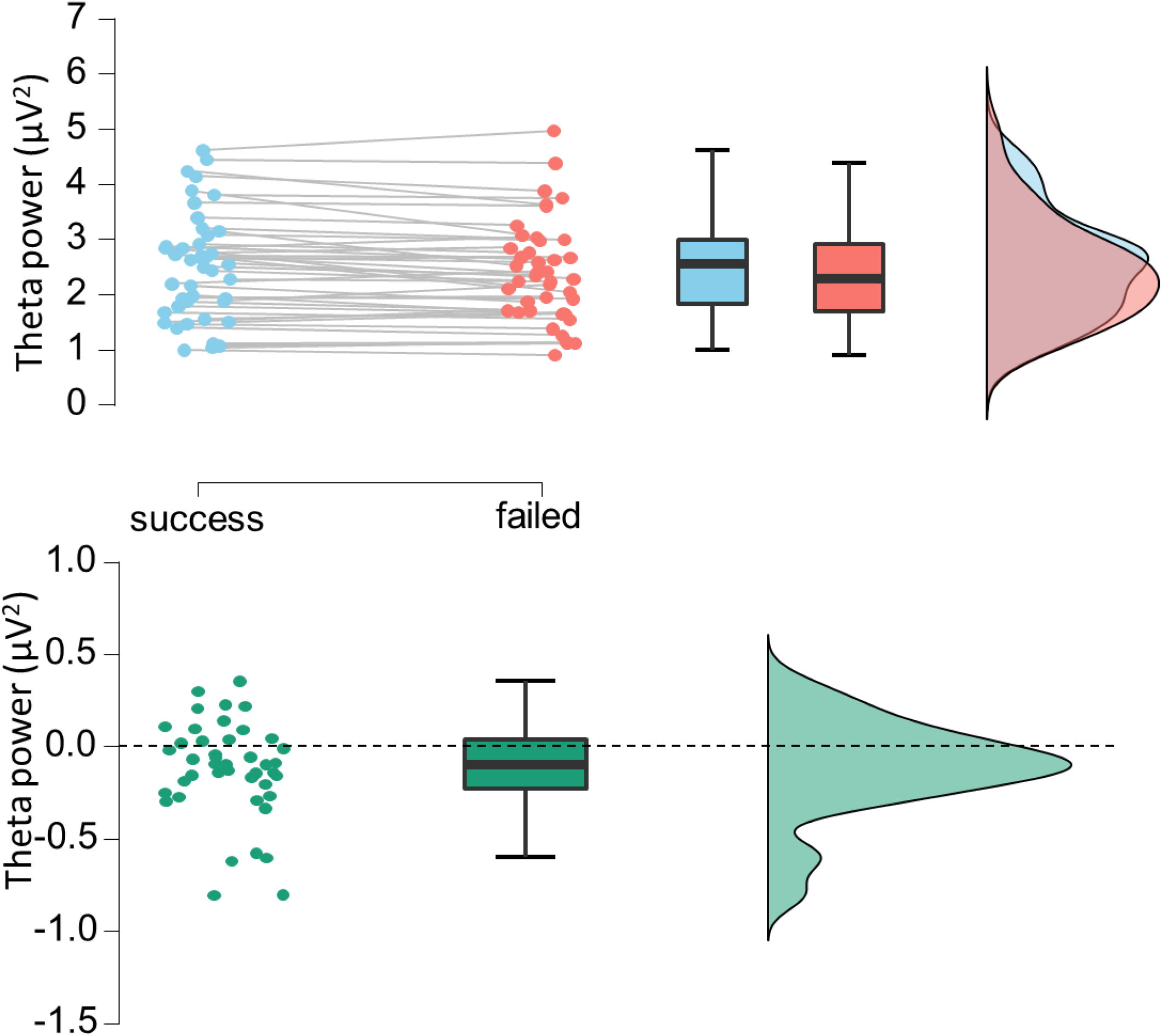
Upper panel: Mean frontal theta preceding failed and successful stop trials. Lower panel: The difference score in mean frontal theta power preceding failed – successful stop trials per subject.

The results from the two-way within-subject ANOVA comparing theta power preceding failed versus successful stop trials in these both conditions are depicted in Table 2. This analyses showed a main effect of stopping performance. Fz theta power preceding failed trials was lower than preceding successful trials (average theta (SDs)=2.39 (0.95) vs. 2.51 (0.9) respectively, collapsed over auditory and visual conditions (see Table 1; Table 2; the difference scores of mean theta power preceding successful – preceding failed trials are depicted in Figure 4). There was no significant difference in theta power between auditory and visual stop-task conditions, nor was there a significant interaction between stop-task condition (Auditory/Visual) and stopping performance (see Table 2).

**Table 2.** model summary of the two-level within subject ANOVA comparing average theta power values preceding both the failed and successful trials for both auditory and visual conditions.

| **Analyses (N=47)** | **F(1,46)** | **$\eta_p^2$** | **p** |
| --- | --- | --- | --- |
| failed/successful | 9.3 | 0.182 | 0.004** |
| auditory/visual | 1.6 | 0.038 | 0.20 |
| failed/successful * auditory/visual | <0.001 | <0.001 | 0.99 |

### Association between SST performance and theta power

The results of the linear regression model are depicted in Table 3. This analysis indicates that mean SSRT was significantly associated with frontal theta power (*B* = −0.26, *t* = −2.63, *p* = 0.009; see Figure 5), indicating that higher theta values are associated with shorter SSRTs. This effect was not significant for go-RT (*B* = 0.12, *t* = 1.32, *p* = 0.19; see Figure 6), nor did the stop-task condition (Auditory/Visual) associate with theta power. Post hoc analyses also indicate that stop-task condition (Auditory/Visual) did not significantly influence the association between SSRT and theta power.

**Table 3.** model summary of the linear regression model estimating the effect of SSRT, go-RT and modality on theta power.

| Analyses | *B* | *t* | *p* |
| --- | --- | --- | --- |
| Mean SSRT | -0.26 | -2.63 | 0.009 ** |
| Mean go-RT | 0.12 | 1.32 | 0.19 |
| Auditory/visual | 0.1412 | 0.74 | 0.46 |

**Figure 5.**
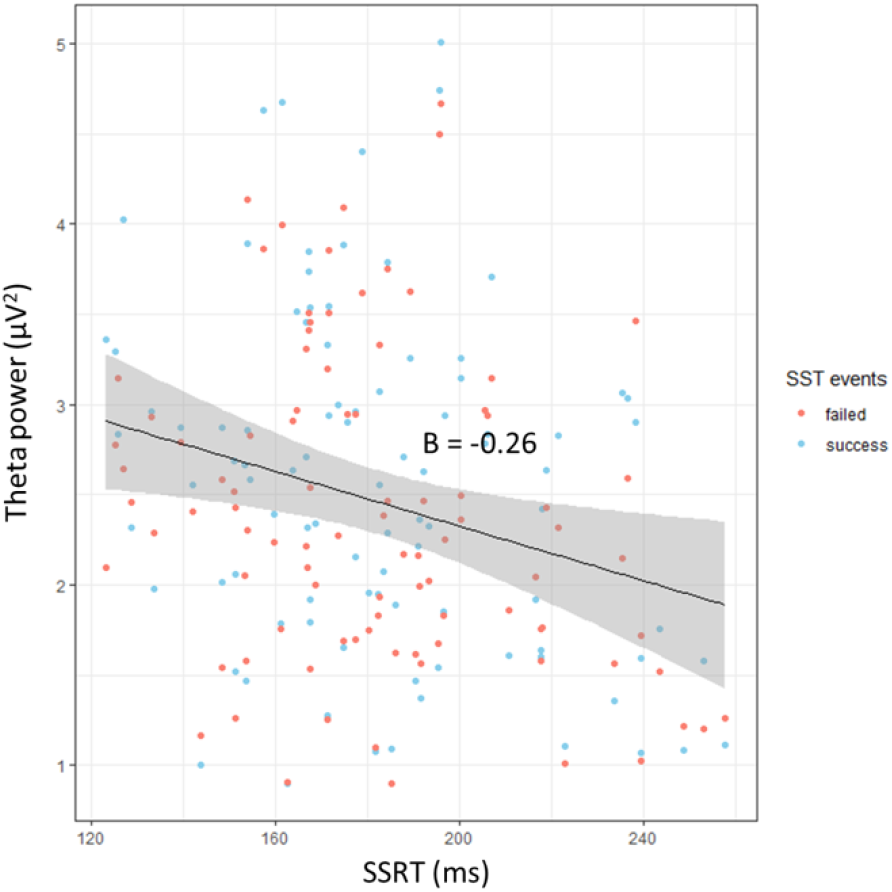
Association between average frontal theta power and SSRT values.

**Figure 6.**
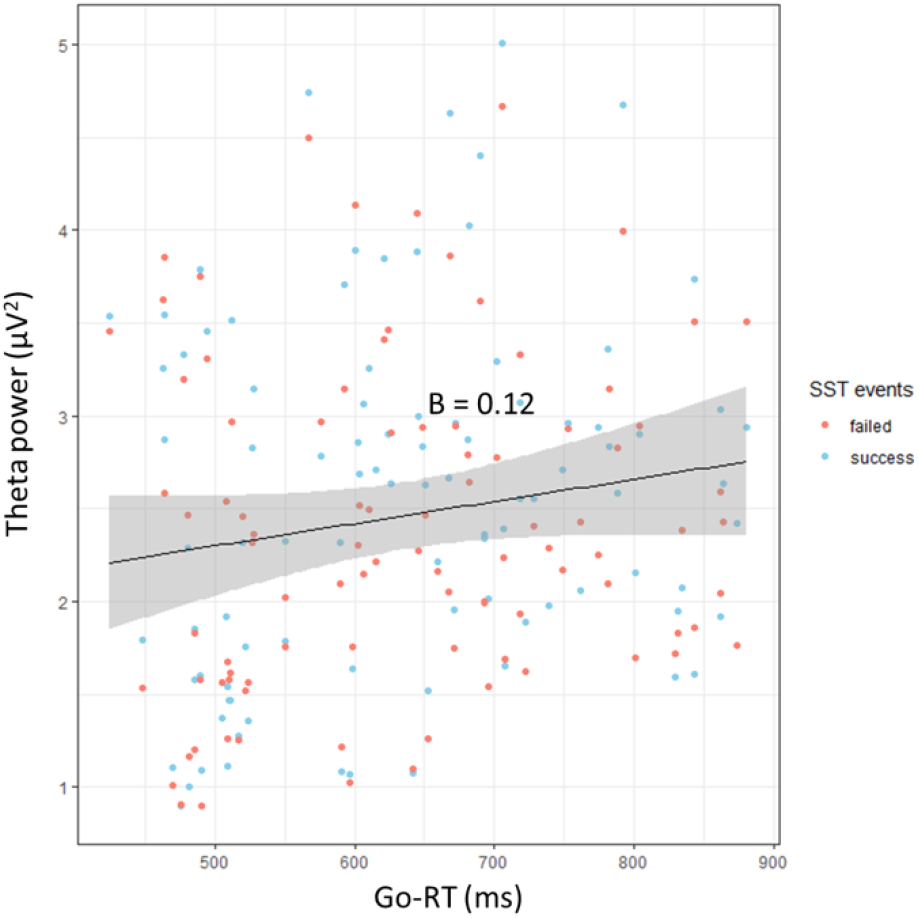
Association between average frontal theta power and go-RT values.

**Figure 6.**
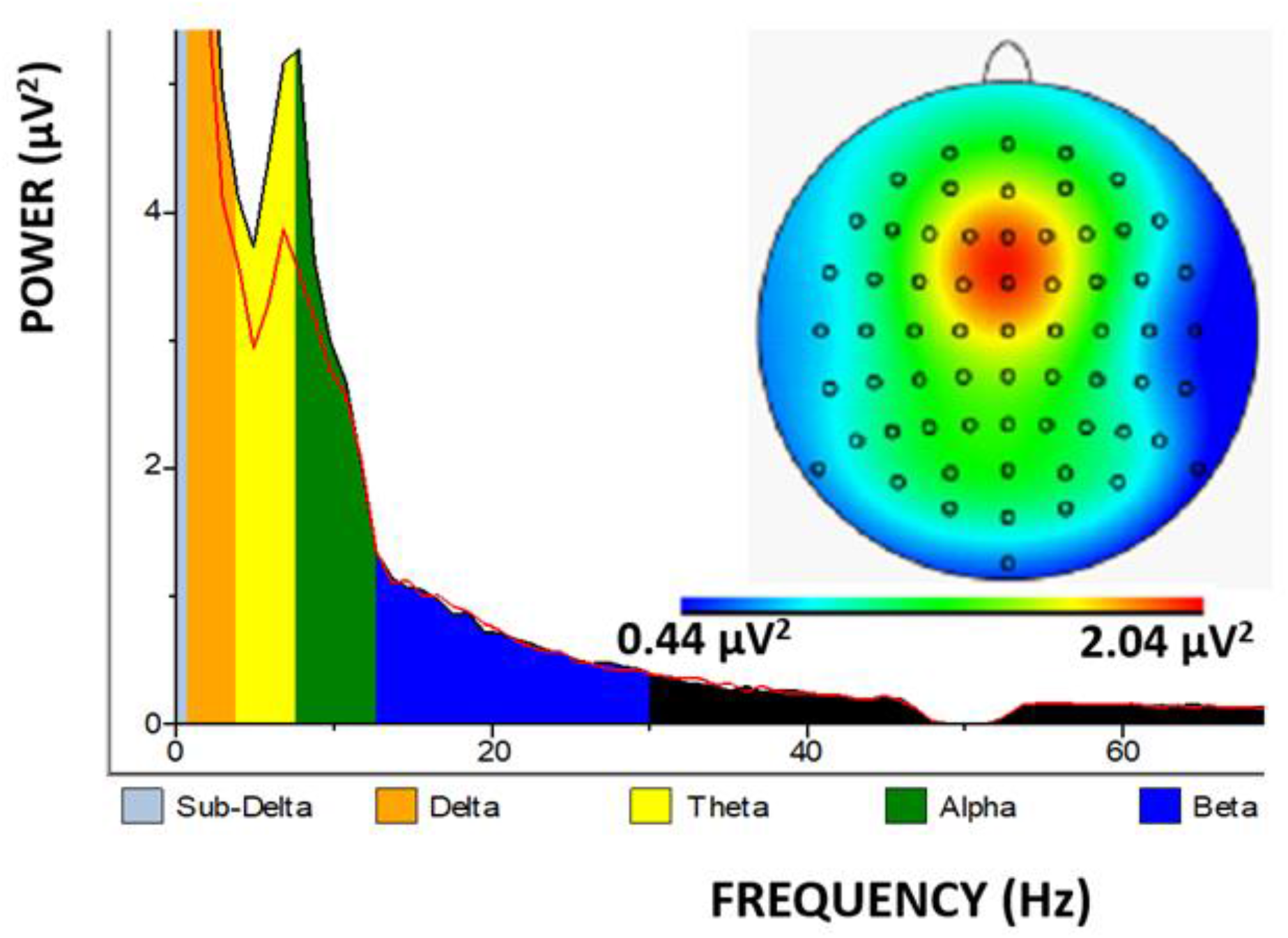
Replication sample: Average power spectra at Fz for long-SOA successful stops and (red line) short-SOA failed stops (black line). Right-insert scalp distribution reflects average across all participants and failed and successful stops.

### Frontal theta power in the replication sample

The results of the replication analyses are depicted in Table 4 and Figure 7. This analysis shows that theta power preceding failed trials was lower than preceding successful trials (means (SDs)=3.59 (2.0); 3.27 (1.9) respectively (*t*(28) = 3.014, *p* < 0.006, d = 0.57)). The secondary analysis indicated that theta power preceding failed trials with a short SOA was lower than preceding successful trials with a long SOA (means (SDs)=3.65 (2.1); 3.15 (2.0); (T(28)=2.63; p<0.014, d = 0.49)).

## Discussion

In this work we investigated whether frontal theta power preceding a stopping event was predictive of stopping success. We demonstrate that in the context of an SST, frontal theta is higher preceding successful stopping events than preceding failed stopping events. We subsequently showed that frontal theta power preceding all stopping events is associated with specifically the SSRT, with a higher theta being indicative of shorter SSRTs. This association was not present for the go-RT across all go trials.

Our within-subject analyses shows that although the mean difference between theta distributions is small, the large majority of our subjects show a decreased mean theta power preceding failed stops. We also show that this is irrespective of the modality of the stop-signal, we observe no difference between conditions with visual versus auditory stop cues.

The explorative analyses in our replication sample further indicate that when we sort our stop events based on difficulty (short vs long SOA), we find a larger mean difference in theta power between failed stop trials with a short SOA (easy) vs successful stop trials with a long SOA (difficult), as compared to the main analyses including all trials. This finding indicates that participants showed higher frontal theta preceding difficult stop trials with successful stops, compared to easy stop trials with failed stops. Note that trial numbers in the easy-difficult comparison were only half of those in the original comparison, which may have rendered statistical significance weaker.

As far as we know, this is the first demonstration that frontal theta is specifically related to stopping success in the context of response inhibition. In particular, our results suggest that frontal theta may be a reflection of proactive response inhibition. More specifically, frontal theta power would be a reflection of the mechanism that proactively facilitates an inhibitory connection between the cortical sensory representation of the to-be-presented-stop-signal on the one hand and the motor system, as postulated in Kenemans (2015).

As mentioned in the introduction, it is important to dissociate this proactive response inhibition from general proactive slowing. A number of studies has shown that when subjects expect something like an odd event, stop-signal or no-go signal, they tend to slow their overall response to the task stimuli (Vink et al., 2014; Zandbelt et al.; 2013; Pas et al., 2017; Pas et al., 2021). However, we demonstrate that frontal theta is not associated with slower go-RTs, which is expected if theta power is associated with general proactive slowing. Our combination of results hence indicate that theta power reflects proactive response inhibition.

In hindsight, the significant correlation between theta power and SSRT seems somewhat surprising, as theta power may be determined by many non-cognitive physiological factors-such as skull thickness and skin conductance. However, previous literature has also shown a relation between theta power and a behavioral measure in the context of motivated learning (Massar, Kenemans & Schutter, 2014). An implication of this is that, had non-cognitive factors been controlled for, the association across individuals between theta and SSRT may have been even stronger.

These results have implications for the research of response inhibition in subjects with ADHD, where response inhibition deficits are one of the most robust cognitive alterations (Lijfijt et al., 2005; Lipszyc & Schachar, 2010). Previous studies have reported no association between go/no-go probability and theta power (Adelhofer et al, 2020), nor differences between ADHD and controls in theta power preceding go or no-go events (Adelhöfer et al., 2021). This may be different when comparing successful versus failed inhibition, and that could help explain the nature of the relation between theta power during cognitive control and ADHD.

An important additional observation in previous literature is that treatment of ADHD patients with methylphenidate largely normalized their stop-signal performance (Overtoom et al., 2009). In addition, global task-related theta power, relative to testing-state values, increases in healthy controls but not in ADHD patients. In turn, methylphenidate restores this increase in ADHD (Skirrow et al., 2015). This would also be consistent with the notion that the deficiencies in response inhibition previously observed in individuals with ADHD might be specifically due to decreased proactive inhibition associated with lower theta power.

In conclusion, to the best of our knowledge, our study is the first to identify a distinct relationship between lower frontal theta power and subsequent stopping failure, highlighting its role in proactive response inhibition. This finding, distinct from proactive slowing, has profound implications for future research in response inhibition and offers new perspectives for understanding and modelling cognitive control mechanisms.

## Author contributions

D. van Rooij: Data acquisition, Formal analysis, Writing

S. van Bijnen: Data acquisition; Review

I. Schutte: Data acquisition; Review

N. Stoep: Methodology; Review.

J.L. Kenemans: Conceptualization; Methodology; Formal analysis; Review

## Notes

### Competing Interest Statement

The authors have declared no competing interest.

